# SLAB: Simultaneous Labeling And Binding affinity prediction for protein-ligand structures

**DOI:** 10.1101/2025.06.03.657720

**Authors:** Aditya Ranganath, Hyojin Kim, Heesung Shim, Jonathan E. Allen

## Abstract

Machine learning models are often used as scoring functions to predict the binding affinity of a protein-ligand complex. These models are trained with limited amounts of data with experimentally measured binding affinity values. A large number of compounds are labeled inactive throughs single-concentration screens without measuring binding affinities. These inactive compounds, along with the active ones, can be used to train binary classification models, while regression models are trained using compounds with binding affinities only. However, the classification and regression tasks are often handled separately, without sharing the learned feature representations. In this paper, we propose a novel model architecture that jointly performs regression and classification objectives, aiming to maximize data utilization and improve predictive performance by leveraging two complementary tasks. In our setup, the regression yields the binding affinity, whereas the classification task yields the label as active or inactive. We demonstrate our method using PDBbind, the standard 3D structure database, as well as a dataset of flavivirus protease compounds with binding affinity data. Our experiments show that the new joint training strategy improves the accuracy of the model, increasing applicability in various practical drug screening scenarios.

## 1 Introduction

The demand for efficient drug screening has surged over the past decade, driven by the emergence of new viral infections and the need for new treatments. This urgency is amplified by the limitations of traditional drug discovery methods, which are time-consuming and resource-intensive. Recent advancements in machine learning (ML) and deep learning (DL) give promising new tools that enable researchers to explore molecular structures with an improved balance between accuracy and efficiency. Deep learning models for binding-affinity prediction, address one of the tasks of assessing the strength of interaction between a protein and a ligand and are critical to accelerating the identification of potential drug leads.

Deep learning models have become widely used as scoring functions for predicting protein-ligand binding affinity. These models leverage molecular structure and various descriptors to learn complex interactions between proteins and ligands. Commonly used descriptors include atomic coordinates, atomic weights, interatomic distances, physiochemical descriptors and molecular fingerprints ^1,2^, which serve as input features for deep learning architectures ^3–7^. By capturing these structural and physicochemical properties, these approaches aim to improve binding affinity prediction accuracy over traditional docking-based methods.

A popular approach that effectively exploits structural information in graph-based representations is the use of *equivariance*. Equivariance is a property of a function or a model where the output changes in a predictable way when the input is transformed. Specifically, a function *f* is equivariant to a transformation *G* if applying *G* to the input and then applying *f* is the same as applying *f* to the input and then applying some corresponding transformation *G*^*′*^ to the output. The most common types of models that use this equivariant information include the SE3 transformer ^8^ and its graphical counterpart E(n) Equivariant Graph Neural Networks (EGNN) ^9^. The EGNN model has been used most recently in predicting protein binding sites by Zhang et al. ^10^, where the authors predict the ligand binding site and the relative direction of the pocket to compute the direction of its nearest ligand atom.

Fusion networks have emerged as a popular approach, where two networks extract different feature modalities from structural data and fuse these features to predict the binding affinity of a protein-ligand pair. One such example is Jones et al. ^11^, where the authors use a Spatial graph CNN (SGNN) and a 3D-Convolutional Neural Network (3D-CNN) ^12,13^. The 3D-CNN voxelizes the 3D space around the protein-ligand pair and extracts the features using a 3D convolutional layer. The SGNN architecture uses the protein-ligand pair as a 2-D graph adjacency matrix representation. These features are then merged and combined to predict the binding affinity of the protein and ligand. These multi-feature learning models showed improved accuracy, but do not scale well on large training sets and as the network becomes too large.

Another multi-head learning approach is GanDTI ^14^, where the authors extract the ligand features using a residual Graph neural network from the SMILES strings and the protein features using an attention network module. The features are then concatenated and fed to an MLP which predicts both the classification of binding inhibition and the binding affinity of the protein-ligand complex structures. One drawback of this approach is that the authors do not use structural information, but rather use feature descriptors to train the models resulting in a lack of information for the model to exploit.

DL approaches also face a challenge with the availability of data. This essentially stems from the lack of ligands and appropriate target protease pairs. To tackle this, researchers often resort to docking methods. Docking methods involve structurally placing the ligand in a protein pocket to analyze their binding energy based on their physical properties. While their predictions are notoriously noisy, they can help generate “hit” compounds without requiring experimentally determined co-crystal structures, which are expensive to generate. Docking and virtual screening pipelines use scoring functions to rank putative poses of a particular ligand at a binding site. The fastest and most widely used method to predict binding affinity is Auto-Dock Vina ^15,16^. AutoDock Vina uses an AD4 scoring function, which is a physics-based model with van der Waals, electrostatic, and directional hydrogen-bond potentials derived from an early version of the AMBER force field. Thus, when the number of experimental data is limited, docking provides an alternative resource of simulation data to understand the binding affinities of compounds and targets.

Existing approaches essentially pose the problem of binding affinity prediction in two ways binding affinity regression problem or activity classification problem. A key limitation of these existing methods is the failure to incorporate features and information from both from the binding affinity values and their activity classification. For example, in a typical experimental compound library screen against a protein target, the vast majority of the tested ligands will show no binding activity. Yet, a docking program can attempt to virtually place the ligand in a target pocket and attempt to score the binding affinity. Similarly, for previous structure-based deep learning methods, the common training set for learning protein-ligand interactions comes from PDBbind, where all of the examples are from ligands that show some binding activity. However, these examples do not contain any examples of ligands which do not show any binding affinity.

To address this type of problem, DL approaches often incorporate a multi-task loss ^17,18^ which motivates a deep learning model to learn multiple tasks in conjunction. Multi-task loss in deep learning is used when a model is trained to perform multiple tasks simultaneously. Instead of optimizing for just one objective, the model learns to minimize a combination of losses from multiple tasks. These kind of multi-task losses can improve the robustness of the model owing to their shared representations. ^19^ The loss function for a multi-task loss is often represented as

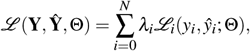

where **Y** = *y*_0_ … *y*_*N*_ are the different labels and **Ŷ** = *ŷ*_0_ … *ŷ*_*N*_ are its corresponding predictions by the model and *λ*_*i*_ *∈* ℝ is a scalar weighing constant. An important observation in the above equation is that the parameters Θ of the model are shared by all the objective loss functions. In other words, the parameter space Θ learns from all the existing loss functions applied to it.

A multi-objective loss function can enhance prediction accuracy by refining the learning process and improving generalization. Instead of treating binding affinity purely as a regression problem, classification helps the model learn distinct patterns between inactive and active binders, reducing noise and improving overall predictive performance. This approach is particularly useful in handling data imbalance and outliers, as binding affinity datasets often exhibit skewed distributions. By defining clear categories, classification prevents extreme values from negatively impacting affinity predictions. Additionally, classification provides a clear decision boundary, making results more interpretable for drug discovery applications and screening away potential inactive candidates. It also contributes to model robustness by preventing overfitting, especially in deep learning approaches like graph neural networks (GNNs) and transformers. When combined with affinity regression, classification strengthens the model’s ability to generalize, leading to more accurate, stable, and interpretable predictions.

A substantial amount of research has been conducted in augmenting the regression loss function using a fixed threshold classification on the regression values ^6,7,20–23^. However, these approaches typically rely on a fixed threshold applied to regression predictions to classify protein-ligand interactions as active or inactive binders, without jointly training on experimental data with inactive binders.

In this paper, we propose a unified framework that integrates both classification and regression, using a novel composite multi-loss scoring function to jointly predict binding affinities and active/inactive labels from protein-ligand structure data. The contributions of our approach are as follows:

1. The proposed approach enables training on both inactive compounds and those with varying binding efficacy by simultaneously optimizing complementary classification and regression objectives.
2. This approach maximizes data utilization by incorporating inactive compounds without binding affinity and compounds with binding affinity, aligning with practical drug screening scenarios.
3. Although we present only a few specific models, the method can be adapted to different network architectures.
4. The framework is lightweight, augmenting the size of the network architecture by additional 40 floating point parameters.

## 2 Method

### 2.1 Problem Statement

For the docking (or co-crystal complex) dataset, we use the complex crystal-structure with the highest binding affinity value (or lowest binding energy value). The data is represented as 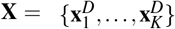, where **x**_*i*_ ∈ *ℛ*^*D*^ represents each data-point’s features with dimensionality *D*. There are two labels for each datapoint - the binding affinity (**y**_*i*_) and its corresponding label (**z**_*i*_). The binding affinity is represented as **Y** = *{***y**_1_,…, **y**_*K*_*}*, where **y**_*i*_ ∈ *ℛ* is a scalar value representing the affinity for **x**_*i*_. For the label, we divide each dataset into two parts - weak binders and strong binders. We set an inactive threshold (IT) for each data point and assign a label **z**_*i*_ to it. Given a threshold IT, the label **z**_*i*_ is defined as

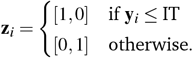

Thus, if the binding affinity value is less than IT, it is labeled as weak, otherwise a strong binder. The main goal is to predict **y**_*i*_ and **z**_*i*_ given **x**_*i*_. Each instance **x**_*i*_ ∈ *ℛ*^*D*^ contains the atomic representation such as the 3D coordinates of the atoms and their atomic features. We describe the data and describe IT in detail in section 2.2 and section 4.

### 2.2 Data

We evaluate our proposed approach on 3 datasets - PDBBind, dengue, and zika (flaviviruses).

#### PDBBind

The PDBbind 2020 dataset ^24^ is a comprehensive collection of protein-ligand complexes with experimentally measured binding affinities, primarily used for training and bench-marking molecular docking, scoring functions, and AI-driven drug discovery models. It is derived from the Protein Data Bank and includes binding data from experimental techniques such as isothermal titration calorimetry, surface plasmon resonance, and enzyme inhibition assays. PDBBind is the collection of experimentally measured binding affinity data (in the form of *K*_*d*_, *K*_*i*_ or IC_50_ values). The dataset is divided into general, refined and core subsets and contains a total of 23, 496 bio-molecular complexes in PDB, which includes protein-ligand (19,443), proteinprotein (2,852), protein-nucleic acid (1,052) and nucleic acid-ligand (149) complexes. It is a fairly balanced dataset where most complex structures have a non-zero normally distributed binding affinity values. The dataset is divided in to general, refined and core datasets arranged in an ascending order of binding resolution quality. The core dataset, or CASF-2016 has the highest quality curation based on resolution and nature of the complexes. The distribution of the binding affinities of the data is illustrated in fig. 2. Each atom feature consists of 3D coordinates (x, y, z), amongst other features such as atom hybridization, number of heavy atom bonds, bond properties, partial charge etc. In addition, each data item has its associated label describing whether its active or inactive.

#### Flavivirus

Flaviviruses are an important group of viruses because they include many significant human pathogens such as Dengue ^25–28^, Zika ^29–31^, West Nile ^32,33^ and Yellow fever ^34,35^ viruses, which are responsible for widespread outbreaks and severe diseases. However, due to the lack of West Nile and yellow fever examples, we choose the Zika (pdbid: 6kk4) ^36^ and dengue (pdbid: 2fom) ^37^ proteases as our flavivirus targets. These targets were retrieved from publicly available protein data bank (PDB) ^38^ database. A total of 2807 ligands were gathered from various publication sources. The ligands, provided in SMILES string format, required pre-processing before further computational analysis. Molecular Operating Environment (MOE) was used for ligand preparation, ensuring proper formal charge assignment and generating 3D energy-minimized conformations of the ligands ^39^. For protein preparation, MOE Protein Quick Prep was utilized to refine structures by adding missing hydrogen atoms, assigning partial charges, optimizing side-chain conformations, and resolving structural inconsistencies such as gaps or missing residues.

Once the structures were retrieved, we perform molecular docking to enrich our dataset. Molecular docking was performed using AutoDock Vina 1.2.0, a widely used docking engine that incorporates improved docking methods and an expanded force field ^16^. The docking workflow was implemented within the Conveyor LC pipeline at Lawrence Livermore National Laboratory (LLNL), which enables high-throughput virtual screening in an automated and parallelized environment ^40^. We generated 20 docking poses complex structure, which were further analyzed for binding affinity and molecular interactions. For the dengue dataset, the protein target was docked with 2303 unique ligands. We choose a random split using the AMPL library. ^41^

These datasets provide us with three unique scenarios - With the PDBBind dataset, we are aware of the exact binding affinity values as these have been experimentally tested and curated. In dengue, we choose a large number of active as well as inactive examples. A ligand is considered as inactive if it shows no inhibitory activity against the protein during single concentration screening. With 20 docking poses for each example, the training subset of the dengue dataset has 1019 (20,380 docking) active examples and 909 (18,180 docking) inactive examples. Similarly, the testing dataset has 266 (5320 docking) active examples and 119 (2380 docking) inactive examples. In the case of Zika, the inactive examples only exist in the training set, not in the testing set, providing a unique scenario of classifying a biased test set. The test set Zika contains 89 (1780 docking) examples, while the train set contains a total of 442 (8840 docking) examples with 379 (7580 docking) active examples and 63 (1260 docking) inactive examples. Thus the performance of the proposed approach will depend on how the inactive and active compounds are identified in the training set to generalize well on the test set. We will explore how the results of the proposed approach changes by introducing inactive complexes in the dataset.

### 2.3 Experimental Setup

In this section, we describe the proposed framework illustrated in fig. 1 in detail. For the loss function, we use the unweighted sum of the cross-entropy (CE) loss and the Mean-squared error (MSE). The CE loss is used for the label classification while the

**Fig. 1.**
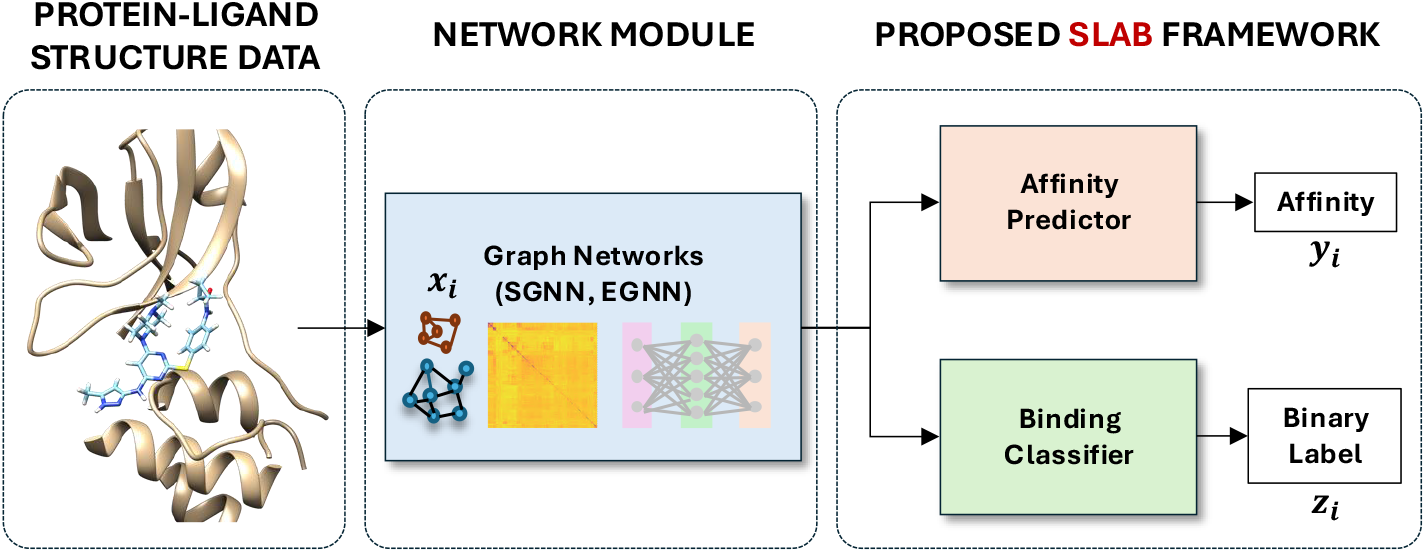
Proposed SLAB framework. The structural data with its corresponding features are fed to a network module (SGNN and EGNN). The affinity predictor module and the label classifier take the output of the network modules to predict the affinity and the label simultaneously.

**Fig. 2.**
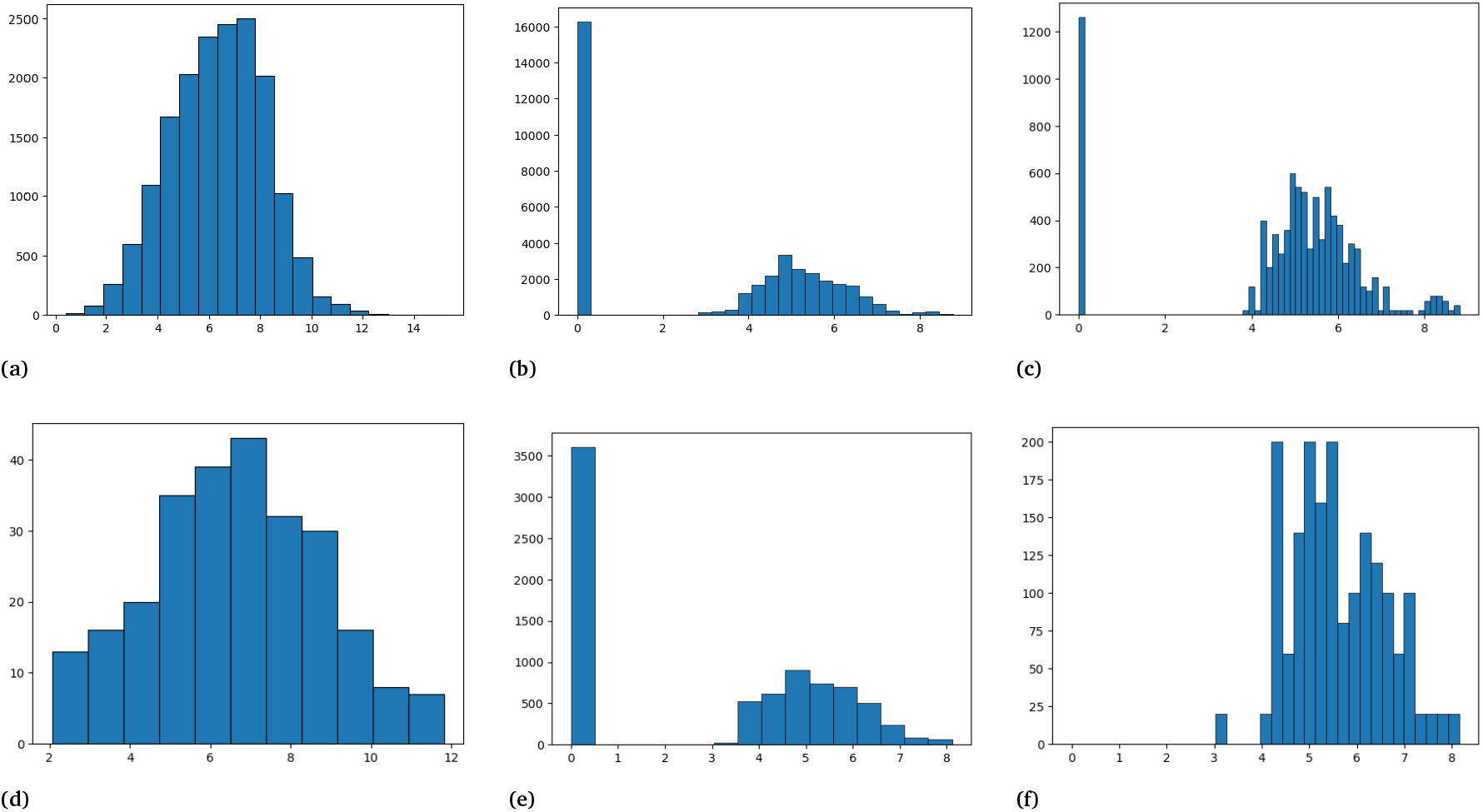
The figure illustrates the distribution of binding affinities from all the datasets in use. fig. (a) and (d) represent the PDBBind train and test set, fig. (b) and (e) presents the dengue train and test set, fig. (c) and (f) present the zika train and test set respectively.

MSE is used for the binding affinity prediction, given by

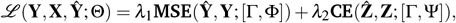

where *ℒ* is the loss function, Θ is the parameters of the network, Φ is parameters of the affinity predictor, Ψ is the parameters of the label classifier and Γ is the parameters of the network module. 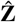 is the predicted labels, *λ*_1 &_ *λ*_2_ are scalars and **Y** is the predicted binding affinity values. We use *λ*_1_ = 1 and *λ*_2_ = 1.

#### Models

We experiment the proposed framework with two very common graphical network modules - EGNN ^9^ and SGNN. We use the Adam ^42^ optimizer to train these neural networks with a learning rate of 1 *×* 10^*−*3^ over 300 epochs. The EGNN has 4 equivariant Graphical convolutional layers with residual connections. The SGNN uses a graph neural network concept, by weighing the bonds between different atoms of the molecule and applying a graphical convolutional layer. We set the covalent threshold of 1.5 Å(cut-off distance for covalent-bonds) and a non-covalent neighbor threshold of 4.5 Å. The features from this layer is then fed to a fully connected layer. The features of both the networks are pooled using the `global_avg_pool and global_avg_pool` library in pytorch-geometric ^43^. The SGNN SLAB architecture has a total of 9323 parameters while the EGNN SLAB has 12569 parameters. We also apply a scheduling algorithm which scales the learning rate by 0.1 every 50 epochs to ensure active training in later epochs. However, we notice that the networks train within the first 100 epochs. The networks were trained on 4 AMD MI300A gpus with an Intel Xeon E5-2695 processing architecture. In case of the PDBBind datasets, we use the general subset for training. For evaluation, we test on the CASF-2016 core dataset. The networks were trained with a batch-size of 32 complexes.

#### Inactivity

For the dengue and Zika datasets, we introduce an inactive assignment (IA) value. For both dengue and Zika, we assign IA = 0 (IA^0^) for all the inactive ligands against the dengue and Zika proteases. Even though the PDBBind dataset does not contain any compounds with a binding affinity value of 0, we maintain the inactive assignment IA^0^. In such cases, instead of drug-screening between active and inactive complexes, the label-classifier behaves like a regularization term ^44^, conditioning the overestimated and the underestimated complexes. In addition to IA, we introduce another value for the dengue and Zika dataset -Inactive Threshold (IT), which we briefly explain in section 2.1. During concentration screenings, low inhibition scores (*y*_*i*_ *<* 3, for example) are deemed inactive owing to their poor inhibitory behavior. Hence, it is safe to consider these as inactive examples, even though they show some inhibitory value in a single concentration screening. We discuss this in more detail in the discussion section (see section 4).

## 3 Results

The results are broadly divided into 3 sections - PDBBind, Dengue, and Zika. The PDBBind data is curated based on the measured binding affinities from multi-concentration measurements. This dataset is used to evaluate performance where ligand-complexes structures are explicitly solved using crystallo-graphic methods (crystal-structures). For our experiments, we treat the general dataset as training data and CASF2016 as our test set. The flavivirus data (Dengue and Zika) reflect more practical cases where there is a combination of single concentration screens and measured activity reported as IC50 with no crystal complexes. We follow from Kim et al. ^45^ and choose the Root-Mean Square Error (RMSE), Mean Absolute Error (MAE), coefficient of determination (**r**^2^), Pearson correlation (R) and Spearman correlation (*ρ*) metrics. We present the results of the proposed SLAB framework using a network module of SGNN ^12^ and EGNN ^9^. Any of these models can be used as the network module to train the network. We choose the SGNN and EGNN as our network module here owing to the ease in training these models.

### PDBBind

The PDBBind data results are shown in table 1. Rows 1 and 3 present the results without using the proposed SLAB framework while rows 2 and 4 present the results using the SLAB framework. We notice that the EGNN-SLAB architecture is able to perform comparably all the other approaches even with one class. In the case of SGNN-SLAB architecture, the model is able to outperform its SGNN alternative in all metrics except RMSE and *r*^2^. Since all compounds in the PDBBind dataset were labeled as active, the predictions of the label classifier were active as well. Hence, we do not present a classification score for this dataset.

**Table 1.** The table shows the results of testing the models on PDBBind2020 CASF-2016. The first section shows the EGNN frame work (rows 1-2), the second section (rows 3-4) shows the SGNN models both with and without the SLAB framework. From these results, it is clear that the proposed SLAB framework with an EGNN network module is able to perform comparably to the other approaches.

| Approach | RMSE | MAE | $r^2$ | Pearson (R) | Spearman ($\rho$) |
| --- | --- | --- | --- | --- | --- |
| EGNN | 1.3113 | 1.0604 | 0.6249 | 0.8040 | 0.7764 |
| EGNN-SLAB | **1.2883** | **1.023** | **0.6380** | **0.8051** | **0.778** |
| SGNN | 1.3950 | 1.12696 | 0.5759 | 0.7726 | 0.7593 |
| SGNN-SLAB | 1.3961 | 1.1115 | 0.5752 | 0.7858 | 0.7681 |

We present the scatter plot of each complex in CASF-2016 dataset in fig. 3. We draw the reader’s attention towards the binding affinity at the higher end (specifically with a binding affinity greater than 10). These examples (fig. 3 (b)) have a higher correlation than the examples where no label (fig. 3 (d)) classifier was applied, thus showing a higher correlation in comparison. This correlation improvement is reflected in data-points with a lower binding affinity as well. Even though the SLAB architecture with the EGNN network outperforms the SGNN network, the relative improvement is comparable. However, the improvement margin of the SLAB architecture is much higher from the regular regression model.

**Fig. 3.**
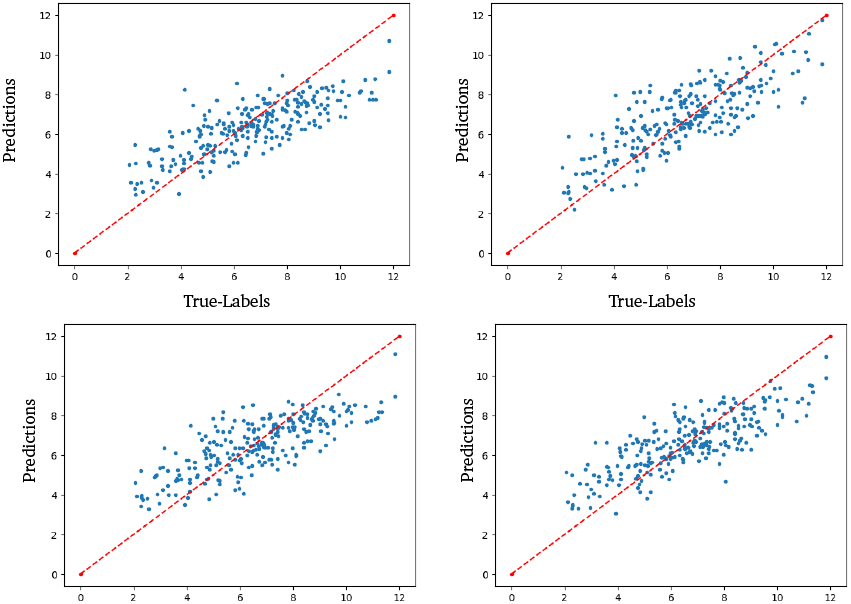
Scatter plots of binding affinity prediction versus experimental value on the PDBBind CASF-2016 dataset. (a) SGNN-SLAB, (b) EGNN-SLAB, (c) SGNN and (d) EGNN. From the figure, and the metrics from table 1, the proposed approach with the EGNN network module performs comparably or better than all other approaches.

### Dengue type-2

Given that the dataset contains 20 docking poses for each compound, we report the results using the average predicted binding affinity across all docking poses. In table 2 we present the outcomes of both the standard regression task and the SLAB approach. With the addition of substantial inactive compounds in the training set, we observe that the SLAB method outperforms significantly across all metrics. Specifically in case of EGNN-SLAB, we notice a significant improvement in all metric values against both of the SGNN models and EGNN without the proposed framework. For more details on the classification accuracy, please refer section 4, experiment IA^0^.

**Table 2.** The table presents the metric values of Dengue type-2. The best values for each metric are highlighted in bold. The superscript ‘avg’ describes the averaging of the binding affinity values over the 20 docking poses. The EGNN-SLAB approach achieves the best performance across all metrics, demonstrating its effectiveness in predicting binding affinities.

| Approach | RMSE | MAE | $r^2$ | Pearson (R) | Spearman ($\rho$) |
| --- | --- | --- | --- | --- | --- |
| EGNN | 1.6527 | 1.2208 | 0.6287 | 0.7944 | 0.7767 |
| EGNN-SLAB | **1.4978** | **1.0315** | **0.6950** | **0.8337** | **0.8136** |
| SGNN | 1.7633 | 1.0867 | 0.5403 | 0.7415 | 0.7021 |
| SGNN-SLAB | 1.6563 | 1.0977 | 0.5944 | 0.7793 | 0.7052 |

### Zika

The results of training and testing the models on the Zika dataset are shown in table 3. The SLAB architecture with an EGNN network outperforms the SGNN and EGNN on all metrics. However, the SGNN performs comparably with and without the SLAB architecture. We notice that in case of Zika, the correlation values (**r**^2^, Pearson (R) and Spearman (*ρ*)) values are much lower when compared with Dengue and PDBBind results. This can be attributed to the high imbalance between the training and test set in terms of the number of active and inactive examples. We also note that all the examples in the Zika testset have been classified as active. Hence we do not report the classification scores for them. For the reader’s convenience, we have provided the scatter plots for Zika in the appendix section.

**Table 3.** The table shows the prediction accuracy of the model on the Zika virus. The bold metrics show the best prediction accuracy among the proposed approaches. The proposed SLAB architecture with the EGNN network module outperforms all existing approaches. The SGNN approach performs comparably with and without the SLAB architecture. For these results, we do not focus on the accuracy since all the labels in the training and testing sets are positive samples. The predicted accuracy for all approaches listed above is 100 %.

| Approach | RMSE | MAE | $r^2$ | Pearson (R) | Spearman ($\rho$) |
| --- | --- | --- | --- | --- | --- |
| EGNN | 1.2282 | 0.9306 | -0.5615 | 0.4302 | 0.5222 |
| EGNN-SLAB | **1.0679** | **0.7882** | **-0.1804** | **0.4509** | **0.5366** |
| SGNN | 1.1532 | 0.8463 | -0.3766 | 0.2991 | 0.3789 |
| SGNN-SLAB | 1.1589 | 0.8055 | -0.3901 | 0.2453 | 0.4157 |

We discuss the results for the Dengue dataset in more detail (along with their scatter plots) in section 4.

## 4. Ablation study and Discussion

From the experiments, it is clear that the proposed approach is able to perform as well or better than single regression task models in all 3 datasets. In the case of the highly curated datasets, the proposed approach performs marginally better. However, it prevents the model from overestimating binding on lower binding affinity compounds, thus reducing the false positive rate. The Dengue datasets are biased for active ligands. Regardless, there is a high number of inactive compounds in the dataset (see fig. 2). In such situations, we might notice that some of the weaker protein-ligand binders get mis-classified, and the performance might degrade ^46^. This can be concluded from the performance improvement indicated in table 2 (see rows 1 and 2). Thus, in the absence of the classification module, the regression module degrades significantly in performance.

The network module plays an important role in the performance of our framework. In our case the EGNN performed well with or without the SLAB framework, followed closely by the SGNN. We explored other model architectures for the network module, however, other models did not yield competitive results due to their memory footprint or inefficiency in training. The EGNN and SGNN are easy to train and easily reproducible, making it an excellent network module for the SLAB framework.

### Inactive Assignment (IA)

Binding affinity values are expressed as pIC_50_, which is calculated on a logarithmic scale using the formula pIC_50_ = *−* log(IC_50_) using molar units. Since a pIC_50_ value of 0 is undefined on a logarithmic scale, assigning such a value to inactive complexes is not meaningful. Hence, we assign a higher binding affinity value of 1 and 2 for the inactive compounds. For example, if the docking results reveal a binding affinity value of 0, we assign these examples with the value of 1 and 2. The results for IA experiments are presented in fig. 4. The scatter plot for this experiment shows a higher correlation and limits the overestimation of weak binders, and improves across all metrics (see table 4 rows 1-3).

**Table 4.** The table presents the results of binding affinity prediction with the inactive values set to 0, 1 and 2 in dengue dataset. In rows 1-3, we present the result of not using the inactive threshold. In rows 4-6, we present the results of using a threshold. From this table, it has been demonstrated that using an inactive threshold can significantly improve across metrics such as RMSE. However, it can affect the classification accuracy.

| Approach | RMSE | MAE | $r^2$ | Pearson (R) | Spearman ($\rho$) | Accuracy (%) |
| --- | --- | --- | --- | --- | --- | --- |
| IA<sup>0</sup> | 1.4978 | 1.0315 | **0.6950** | **0.8337** | **0.8136** | 88.75 |
| IA<sup>1</sup> | 1.4736 | 1.0474 | 0.5312 | 0.7501 | 0.7306 | 85.18 |
| IA<sup>2</sup> | **1.0791** | **0.8167** | 0.6044 | 0.7907 | 0.7545 | 85.16 |
| IT<sup>3</sup> IA<sup>0</sup> | 1.8768 | 1.4226 | 0.4791 | 0.7352 | 0.6984 | 67.02 |
| IT<sup>3</sup> IA<sup>1</sup> | 1.548 | 1.228 | 0.4823 | 0.6976 | 0.7352 | 76.27 |
| IT<sup>3</sup> IA<sup>2</sup> | **1.1009** | **0.8146** | **0.5883** | **0.7559** | **0.7750** | 81.53 |

**Fig. 4.**
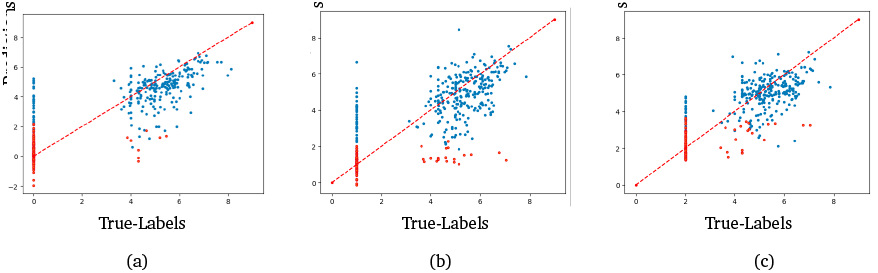
The figure represents the Inactive Assignment (IA) results for the dengue dataset with the inactive assignment set to 0, 1 and 2. fig. (a) represents the results setting the IA^0^, fig. (b) represents the results IA^1^ and fig. (c) represents the results with IA^2^. Blue color represents the complexes predicted as active by the model, while the red color represents the inactive predictions. The figure shows the reassigning the affinity value to a lower bound limits the outliers. This reassignment also boosts the correlation in these cases.

### Inactive Threshold (IT)

As brifly discussed in section 2.3, generally a complex with a low binding affinity value (*<* 3 for example) is considered a weak binder, which is equivalent to calling these complexes as inactive due to a lack of inhibitory behaviour. In such cases, it is safe to consider these examples as inactive and assign the IA to these complexes. To validate this, we define a threshold for the inactive compounds - we set a threshold at pIC_50_ = 3, ensuring that compounds with pIC_50_ values above this threshold are labeled as active binders, while those at or below are labeled as inactive binders. For example, if the inactive threshold is set to 3, all compounds revealed with a binding affinity of lower than 3 are replaced with inactive assignments of 0, 1 or 2. This reassignment to a lower constant inactive assignment also improves the model’s performance by focusing on the higher binding affinity values. This improvement is empirically reflected in the results. Our SLAB architecture is trained based on these labels, with all other network parameters remaining unchanged. Results are shown in table 4 and the scatter-plot for this experiment is in fig. 4. fig. 4 shows the results of setting the binding affinity to 2 for the labels of the inactive examples. In this study, we explore setting a strong threshold for the binding affinities to 3. This means, all the complexes with a binding affinity less than or equal to 3 are set to a binding affinity of 0, 1 or 2. Model accuracy is shown in fig. 5.

**Fig. 5.**
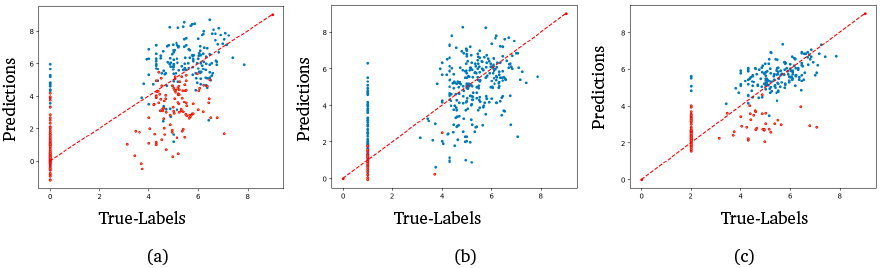
Inactive threshold results for dengue dataset. The threshold value for the compound structures is IT^3^. The points marked in blue are examples predicted as active by the classifying module, while the ones marked in red were labeled inactive by the classifier module. fig. (a) represents IA^1^, (b) represents IA^2^ and (c) represents IA^3^

It is clear from the figure that the proposed approach with a threshold and setting a higher inactive assignment value yields significantly better RMSE values. However, we do notice that the classification accuracy in case of the IT results are compromised. With a lower IA (IT^3^IA^0^ and IT^3^IA^1^), the classification accuracy is much less when compared with IT^3^IA^2^. In case of the IT^3^IA^2^, the experiment yielded the best RMSE value of all the experiments. This experiment yielded the best metrics across all the IT experiments.

### Precision-Recall

fig. 6 shows the precision-recall curve when the model is evaluated as an active/inactive classifier with the binding affinity threshold for actives is set to 3. The *y*-axis is precision, given by 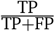 where TP is the True-positive rate and FP is the false positive rate. The *x*-axis is the recall, given by 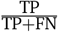, where FN is the False negative rate. The trade-off between correctly identifying positive cases and incorrectly classifying negatives is shown when adjusting the model’s prediction threshold. The Area Under the Curve (AUC) quantifies overall model performance, with values closer to 1 indicating better discrimination.

**Fig. 6.**
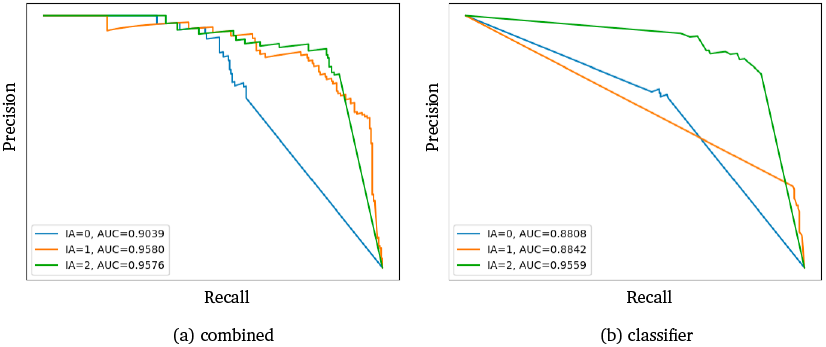
Precision-Recall Curve comparing model performance on dengue dataset. The curve illustrates the trade-off between True Negatives and False Positive Rate (1-Specificity). The *x*-axis and *y*-axis range between [0, 1]. A higher area under the curve (AUC) indicates better classification performance. Fig. (a), represents the curve if both the regression and classification modules are used, while fig. (b) represents the results only with the classification module.We notice that the classification AUC is significantly improved by using both the classification module and the regression module over using just the classification module.

fig. 6 shows performance when using the classifier first, and where the classifier predicts the ligand to be active, the regression value is used at varying thresholds to re-classify the ligand as an active. The model was compared with the baseline of using only the classification module to classify ligands.

We present the results for the threshold results in fig. 6. From this figure it is clear that having a higher lower-bound with a threshold increases the AUC, therefore indicating better discrimination.

### Comparative performance

To show the improvement of combined classification and regression modules against the classifier, we present the results with IA^1^ in fig. 7 with a threshold of IT^3^. We choose this result owing to the improvement from using just the classifier i From these results, it is clear that using both the classifier and regression module is significantly better than just using the classifier.

**Fig. 7.**
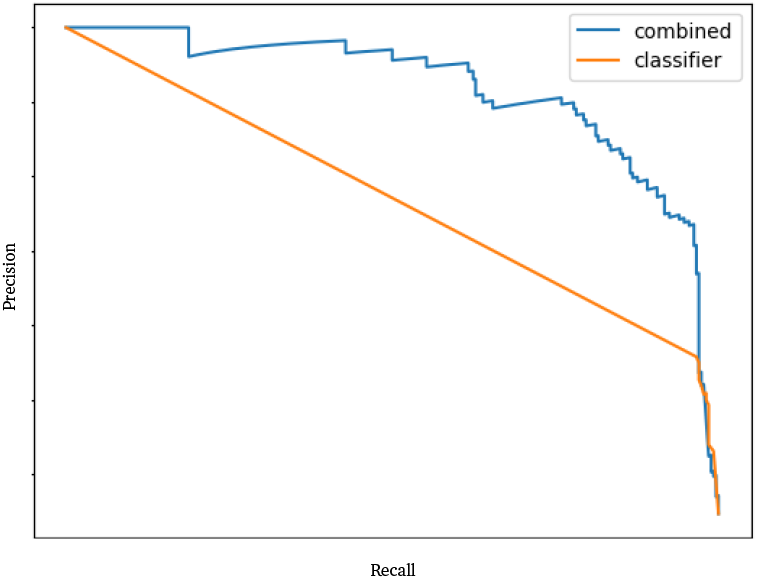
Binding affinity precision-recall curves for dengue dataset with IA = 1. The orange line represents using just the classifier, while the blue line represents using both the classifier and the regressor. From the graph, it is clear that using both the classification module and the regression module a higher AUC than just using the classifier.

## 5 Conclusion

We present a new training framework to incorporate classification into a regression task. Through experimentation, we empirically show that the proposed SLAB approach outperforms existing regression techniques. Further experimentation using a inactive threshold and inactive assignment showed significant improvement in binding affinity value predictions.

We do notice a drop in classification performance with a low inactivity assignment and an inactive threshold value. However, the margins improved as we increased the inactivity assignment values for those thresholds. In addition, the precision-recall curves have clearly illustrated that using the classification module in conjunction with the regression module has yielded in a better AUC when compared against the classifier.

## 6 Acknowledgment

This work was performed under the auspices of the U.S. Department of Energy by Lawrence Livermore National Laboratory under contract DE-AC52-07NA27344 with IM release number LLNL-JRNL-2004933. Lawrence Livermore National Security, LLC. This work was funded by the Defense Threat Reduction Agency (DTRA), HDTRA1242044, HDTRA1036045 and LLNL Laboratory Directed Research and Development 24-SI-006. Computing support for this work came from the Lawrence Livermore National Laboratory (LLNL) Institutional Computing Grand Challenge program.

