## Supplemental figure 8 and figure 9 for "SLAB: Simultaneous Labeling And Binding affinity prediction for protein-ligand structures"

PAPER

Cite this: DOI: 00.0000/xxxxxxxxxx

Received Date
Accepted Date

DOI: 00.0000/xxxxxxxxxx

Machine learning models are often used as scoring functions to predict the binding affinity of a protein-ligand complex. These models are trained with limited amounts of data with experimentally measured binding affinity values. A large number of compounds are labeled inactive through single-concentration screens without measuring binding affinities. These inactive compounds, along with the active ones, can be used to train binary classification models, while regression models are trained using compounds with binding affinities only. However, the classification and regression tasks are often handled separately, without sharing the learned feature representations. In this paper, we propose a novel model architecture that jointly performs regression and classification objectives, aiming to maximize data utilization and improve predictive performance by leveraging two complementary tasks. In our setup, the regression yields the binding affinity, whereas the classification task yields the label as active or inactive. We demonstrate our method using PDBbind, the standard 3D structure database, as well as a dataset of flavivirus protease compounds with binding affinity data. Our experiments show that the new joint training strategy improves the accuracy of the model, increasing applicability in various practical drug screening scenarios.

### Notes and references

### A Results on dengue and zika

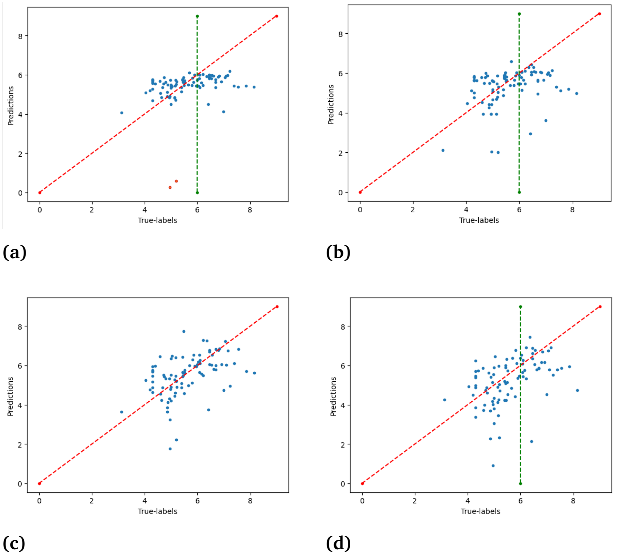


Fig. 1 The figure presents the scatter plots of Binding affinity prediction for zika. (a) represents the SGNN-SLAB results, (b) SGNN results, (c) represents EGNN-SLAB results and (d) represents the EGNN results. From the figure, and the metrics from table 1, the proposed approach with the EGNN network module outperforms the all other approaches.

[a] *Center for Applied Scientific Computing, Lawrence Livermore National Laboratory, 7000 East Avenue, Livermore, California;*
[b] *Biosciences and Biotechnology Division, Lawrence Livermore National Laboratory, 7000 East Avenue, Livermore, California, 94550*
[c] *Global Security, Lawrence Livermore National Laboratory, 7000 East Avenue, Livermore, California, 94550*

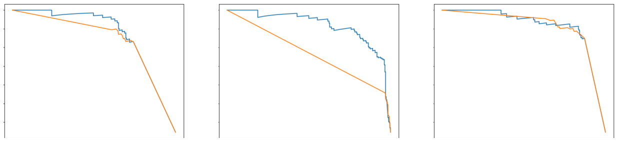

Fig. 2 Binding affinity precision recall curves for different Inactive assignments.
